# Robotic light touch assists human balance control during maximum forward reaching

**DOI:** 10.1101/848432

**Authors:** Leif Johannsen, Karna Potwar, Matteo Saveriano, Satoshi Endo, Dongheui Lee

## Abstract

**Objective:** We investigated how light interpersonal touch (IPT) provided by a robotic system supports human individuals performing a challenging balance task compared to IPT provided by a human partner.

**Background:** IPT augments the control of body balance in contact receivers without a provision of mechanical body weight support. The nature of the processes governing the social haptic interaction, whether they are predominantly reactive or predictive, is uncertain.

**Method:** Ten healthy adult individuals performed maximum forward reaching (MFR) without visual feedback while standing upright. We evaluated their control of reaching behaviour and of body balance during IPT provided by either another human individual or by a robotic system in two alternative control modes (reactive vs predictive).

**Results:** Changes in reaching behaviour under the robotic IPT, such as lower speed and straighter direction were linked to reduced body sway. MFR of the contact receiver was influenced by the robotic control mode such as that a predictive mode reduced movement variability and increased postural stability to a greater extend in comparison to human IPT. The effects of the reactive robotic system, however, more closely resembled the effects of IPT provided by human contact provider.

**Conclusion:** The robotic IPT system was as supportive as human IPT. Robotic IPT seemed to afford more specific adjustments, such as trading reduced speed for increased accuracy, to meet the intrinsic demands and constraints of the robotic system. Possibly, IPT provided by a human contact provider reflected reactive interpersonal postural coordination more similar to the robotic system’s follower mode.

**Précis:** Interpersonal touch support by a robotic system was evaluated against support provided a human partner during maximum forward reaching.

Human contact receivers showed comparable benefits in their reaching postural performance between the support conditions.

Coordination with the robotic system, nevertheless, afforded specific adaptations in the reaching behaviour.

## Introduction

If robotic systems are envisaged as the solution to future shortages in clinical staff and caregivers for the purpose of augmenting of patients’ mobility by a provision of balance support, they must show a responsiveness to the social constraints and demands, which govern any routine physical interaction between a patient and a human carer. From a scientific and engineering point of view, therefore, the principles of human-human interactions during physical interactions need to be extracted and evaluated in terms of their transferability to human-robot interactions as exoskeletal approaches may be unsuitable for frail individuals due the weight added to the body. In physical rehabilitation, caregivers and therapists routinely provide physical assistance to balance-impaired individuals during postural mobilization and transfer maneuvres. In order to prevent long-term habitual dependency of a patient on external balance aids and other forms of support, a therapist needs to be adopt an optimum level of postural assistance that maximizes a patient’s movement autonomy (‘assist-as-needed’). One possible approach is the provision of delibrerately light interpersonal touch (IPT) by a caregiver, which can be used to reduce body sway in quiet standing in neurological patients with impaired postural stability when applied to patients’ backs (Johannsen, McKenzie, Brown, Redfern, & Wing, 2017). In such an interpersonal postural context, the contact receivers (CR) experiences haptic contact passively with little or no possibility to influence the interaction due to their greater motion-task constraints compared to those of the contact provider (CP). Not only the movement degrees of freedom available to each individual during IPT, but also the relative postural stability of both partners determines the strength of the IPC and the individiual benefit of IPT, with more enhanced postural stability in the intrincically less stable person (Johannsen, Wing, & Hatzitaki, 2012).

To explore the interdependencies between CR and CP during IPT in more detail, we evaluated performance in maximum forward reaching (MFR) with and without light IPT applied to the ulnar side of the wrist of blindfolded CR’s extended arm intended to provide a social haptic cue and impose social coordinative constraints on both the CR and the CP (Steinl & Johannsen, 2017). Interestingly, IPT reduced sway more effectively when the CP had the eyes closed and their perception of CR’s motion was based on haptic feedback alone. In contrast, IPT with open eyes did not result in reduced sway compared with a condition in which IPT was not provided (Steinl & Johannsen, 2017). We speculated, therefore, that minimization of the interaction forces and their variability at the contact location during IPT acts as an implicit task constraint and shared goal between both partners (Knoblich & Jordan, 2003). This goal might afford predictive sway control in each individual and consequently led to in-phase interpersonal postural coordination with an average zero lag but also minimization of the variability of the interaction force (Johannsen, Guzman-Garcia, & Wing, 2009; Johannsen et al., 2012).

In the present study, we intended to contrast the effects of human IPT (hIPT) on CR’s postural performance against the effects of two different modes of robotic IPT (rIPT) and expected specific costs and benefits on body sway and postural performance due to the robotic response modes. Similar to hIPT, rIPT was applied in a “fingertip touch” fashion to CR’s wrist without any mechanical coupling or weight support. The robotic system either followed a participant reactively or predicted a participant’s movement trajectory. As the coupling between two humans with IPT in terms of the interaction forces is intrinsically more noisy due to each individual’s motion dynamics and response delays, we expected that a predictive mode of the robotic system would result in a less noisy haptic coupling and therefore enhance performance in the MFR task, such as greater reaching distance with less body sway. In addition, the reactive mode of the robot was supposed to be advantageous over hIPT due to the fixed response delay, which would enable participants to extract own movement-related information from the interaction forces for balance control.

## Methods

### Participants

We tested 10 healthy young adults (average age=28.5, SD 3.35 years, 3 females and 7 males) as contact receivers (CR) performing a maximum forward reaching (MFR) task. Participants were not affected by any neurological or orthopedic indications. Participants were recruited as an opportunity sample from students of the university. The study was approved by the ethical committee of the medical faculty of the TU Munich and all participants gave written informed consent.

### Equipment and experimental procedure

One healthy adult, male contact provider (CP) applied the IPT to the wrists of the contact receivers (CR). The CR stood blindfolded on a force plate (Bertec 4060, Columbus, OH, USA; 500 Hz) in bipedal stance performing the MFR task. CR was always instructed to reach as far forward as possible by bending the torso but not the knees. Before the start of a trial, CR was instructed to stand in a relaxed manner, the right arm extended at shoulder height to reach horizontally above a height-adjusted table. After the start of a trial, CR was instructed to remain static for at least 5 seconds (baseline) until an auditory signal cued the start of the MFR task (Fig. 1a). During IPT, CP stood facing orthogonally to CR in bipedal stance between CR’s force plate and the table, parallel to the reaching direction. CP provided IPT with the right extended arm by lightly contacting the wrist at its ulnar side of the CR. During IPT, CP kept the eyes open to receive visual cues of a CR’s motion as would the robotic systems by optical motion tracking. During the robotic IPT conditions, a single KUKA LWR4+ manipulator (Augsburg, Germany) served as CP. The CP kept light contact with CR’s ulnar side of the wrist. CR’s body sway was determined in terms of the anteroposterior (AP) and mediolateral (ML) components of the Center of Pressure (CoP), as derived from the six components of the ground reaction forces and moments.

**Figure 1.**
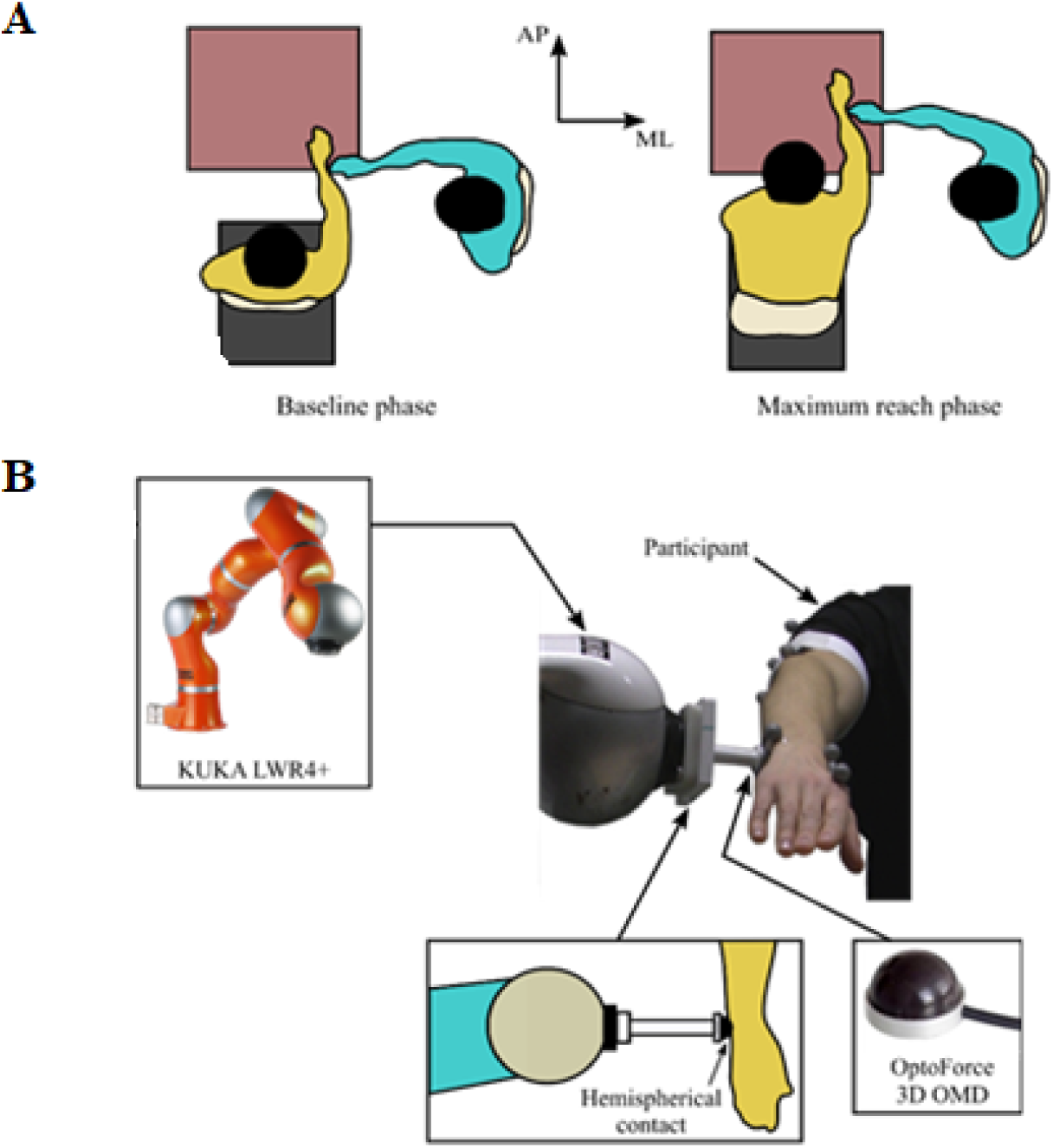
Experimental setup. (A) Execution of the maximum forward reach task with human interpersonal touch (hIPT) support. (B) Robotic IPT without mechanical coupling in hybrid force-position control.

In in the human-robot interaction conditions, the CR’s wrist was tracked by the end effector of the robotic system without any mechanical coupling (Fig. 1b). The robotic system provided contact via a hemispherical rubber pad attached to a force sensor (OptoForce 3D OMD, OnRobot. Odense, Denmark; 500 Hz) on the end-effector, which kept the relative orthogonal distance constant. The force sensor was used to measure force at the contact location. The CR’s wrist position, required to control the robotic system, was measured by an optoelectronic motion capture system (OptiTrack, NaturalPoint, Corvallis, OR, USA; 100 Hz). To provide nearly the same feeling for the CR in both touch conditions, the CP was wearing a thin rubber glove to provide similar tactile sensation to the case of rIPT where the end effector of the robot had a rubber surface (Fig. 1b). Participants’ movements of the right hand were tracked with a marker-based optical motion capture system by placing three reflective markers on the right hand (one on the caput ulnae/processus styloideus radii/basis and two on the ossa metacarpi). Tracked hand position was sent to the robot to control the robots’ movements but also recorded to calculate reaching distance in the MFR end-state. The robotic control scheme required high control frequencies to avoid unstable behaviors (Siciliano, Sciavicco, Villani, & Oriolo, 2009). For this reason, the robot was controlled at 500 Hz. Interaction forces were measured at the same frequency of 500 Hz, while the CR’s hand was tracked at 100 Hz. Hence, it was necessary to up-sample the motion tracking system to match the robot control frequency.

This experiment contrasted three modes of IPT provision: hIPT, robotic light interpersonal touch with reactive following of the participant’s movements (rIPTfollow), robotic light interpersonal touch with anticipation of the participant’s movements (rIPTanticip). Robotic IPTfollow and rIPTanticip were both provided through an artificial “finger” with optical tracking of the CR’s wrist and control of the contacting force. The three IPT conditions were assessed in blocks of 5 trials. The order of the blocked conditions was fully randomized, and each single trial lasted 20 s. Out of a total of 150 trials, 11 trials failed to track the CR’s hand and are excluded from the analysis.

### Data reduction

All data post-processing was conducted in Matlab 2016b (Mathworks, Natick, MA, USA). Kinematic and force-torque sensor data were spline-interpolated to 600 Hz and subsequently merged with the force plate recordings. The data was smoothed using a generic dual-pass, 4^th^ order Butterworth low-pass filter with a cut-off frequency of 10 Hz. CoP and marker data were differentiated to yield velocity. Subsequently, trials were segmented into three phases of the MFR (baseline phase, reaching phase, and MFR end-state) based on the AP position of CR’s wrist marker as described in Steinl and Johannsen (Steinl & Johannsen, 2017) (Fig. 2).

**Figure 2.**
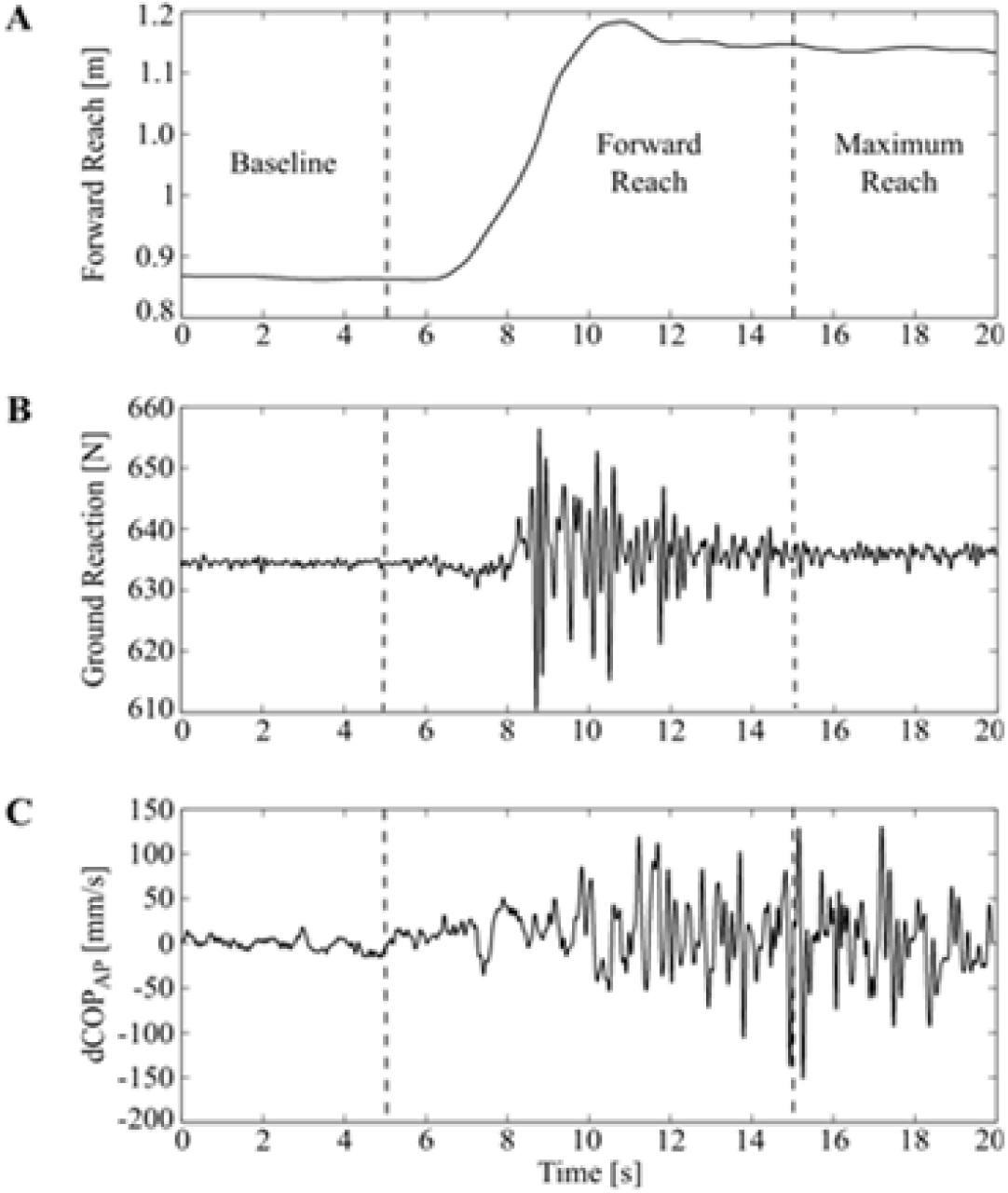
Typical profiles of the kinematic and dynamic variables. (A) Forward reaching of the hand marker divided into three phases. (B) Ground Reaction Force in the vertical direction. (C) Centre-of-Pressure velocity (dCoP) in the anteroposterior (AP) direction.

To investigate the effects of IPT on human CR’s postural performance during the maximum forward reaching (MFR) task, MFR amplitude in the horizontal plane was determined from the difference of the wrist’s average position in the baseline phase and in the MFR end-state. The angular deviation of a straight line connecting these two positions from the AP axis, the path length and normalized path length (path length/amplitude) of the reaching trajectory, the average and summed as well as the standard deviation of the orthogonal deviation of the trajectory from a straight line and the average and peak velocity of the wrist during the reaching phase were extracted. Body sway in the baseline, reaching phase as well as in the MFR end-state was extracted as the standard deviation of the COP velocity (SD dCoP) in AP and ML directions. In order to quantify the efficiency of balance control during the MFR reaching phase and evaluate a potential speed-accuracy tradeoff, we calculated an Index of Performance (IoP) for both sway directions based on a modification of Fitts and Peterson’s index (Bootsma, Fernandez, & Mottet, 2004; Fitts & Peterson, 1964). Our IoP associated the time for the reaching movement (MT) with the difficulty of the MFR (IoD) in terms of the achieved maximum amplitude (A) and the variability of sway in the reaching phase (S) as the effective precision constraint:

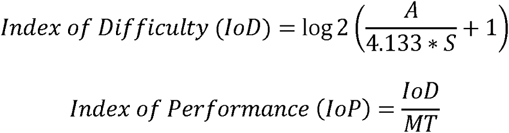

The unit of the IoP is bit/s and thus expresses the informational “throughput” of a participant during the movement.

### Statistical analysis

SPSS version 23 (IBM, Armonk. NY, USA) was used for statistical analysis. All outcome parameters were log-linearized before statistical analysis to approximate normal distribution. A linear mixed model with IPT condition as within-subject factor including participant as random effect was applied using maximum likelihood estimation. To test for statistical significance, an alpha level of 0.05 was used and post-hoc comparisons were computed as required to distinguish between IPT conditions.

### Robotic control

Both the robot end-effector position and the interaction force were actively controlled using a hybrid force-position controller, which was based on the prediction of the CR’s wrist motion. A Linear Kalman Filter (LKF) (Kalman, 1960) with a constant velocity model was exploited to generate a reference for the participant’s wrist trajectory. A constant velocity LKF assumes that the motion is generated by the discrete linear system

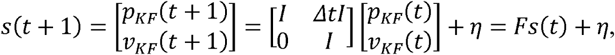

where the state vector *s*(*t*) contains the Kalman-estimated wrist position *p*_*KF*_(*t*) and velocity *v*_*KF*_(*t*), *I* is an identity matrix, Δ(*t*) is the sampling time, and *η* is an additive Gaussian noise. The LKF predicts the next state *s*_*KF*_(*t*) = [*p*_*KF*_(*t*)^T^ *v*_*KF*_(*t*)^T^]^T^ = *Fs*(*t* −1) + *y*(*t*), where the correction term *y*(*t*) is computed as in [56] and it depends on the measured wrist position. In our setup, the correction term was set to *y*(*t*) = 0 until a new measure of the wrist position was available. In this way, the predicted position *p*_*KF*_(*t*) was generated at 500 Hz and used to control the robotic system. The LKF was exploited to realize two different robotic modes, i.e. the robotic follower and the robotic anticipatory modes. More specifically, in the rIPTfollow mode the robot passively followed the wrist motion while providing a light touch. To implement a passive follower, the position *p*_*KF*_(*t*) (Position Error: rIPTfollow AP – 0.010218m, ML – 0.004994 m) (Fig. 3b) predicted by the LFK at the actual time instant t was used to generate the control command described in the previous section. In this way, the robotic system followed the wrist position with one sample delay (10 ms). In the rIPTanticip mode, the robot predicted the future wrist position to lead the motion while providing a light touch. To realize the leading mode, the LKF was exploited to make a one-step prediction of the wrist position. In particular, the predicted future position *p*_*KF*_(*t* +1) = F *p*_*KF*_(*t*) (Position error: rIPTanticip AP – 0.012256, ML – 0.007164 m) (Fig. 3a) was used to generate the control command. In this way, the robot was anticipating the human motion by one sample (10 ms), thereby leading the movement execution.

**Figure 3.**
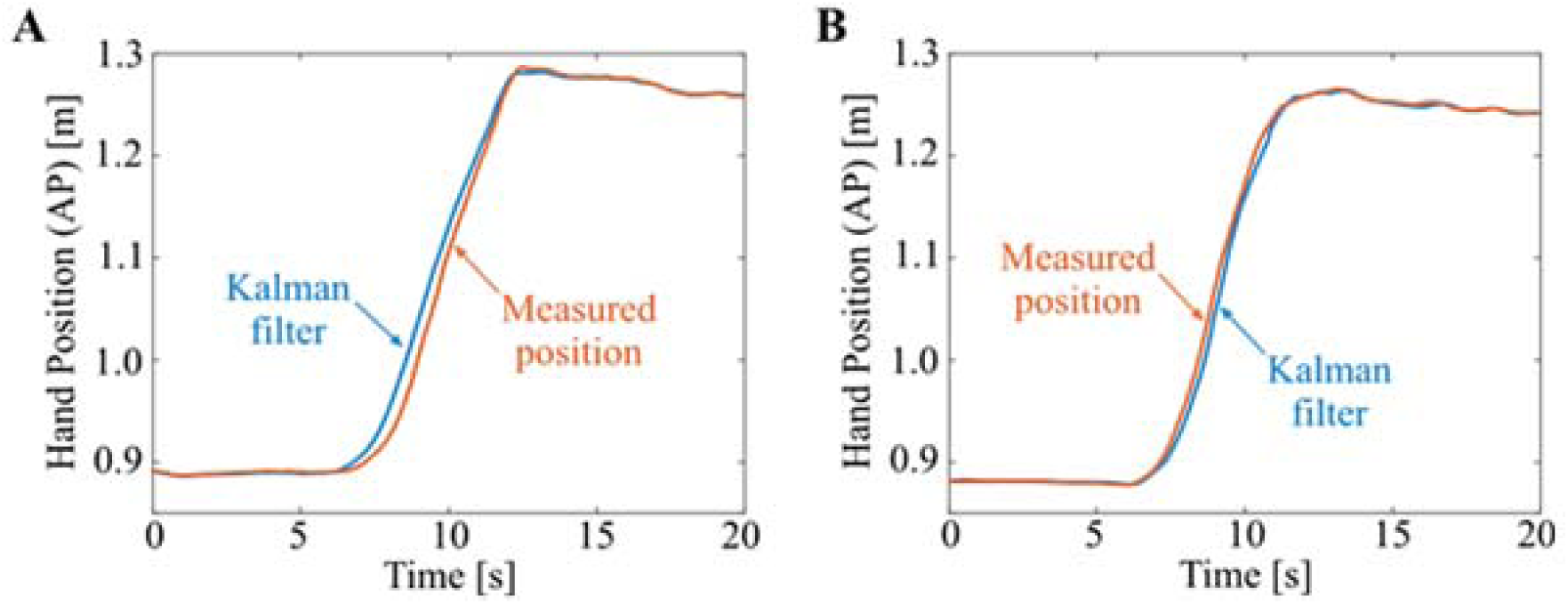
Kalman filtered hand position during maximum forward reaching (MFR). (A) Predicted and measured hand position during MFR for anticipatory robotic interpersonal touch (rIPT) in the anteroposterior (AP) direction. (B) Estimated and measured hand position during MFR for rIPT in follower mode in the AP direction.

During the MFR task, the robotic system provided a light touch along the contact directions, while predicting and following (or predicting) the participant’s right wrist trajectory in the AP direction. The robotic system was controlled to exert a maximum of 1 N force along the ML and vertical directions (force-controlled directions), while tracking the hand motion along the AP axis (position-controlled direction). The force *f*_*m*_ = [*f*_*m,x*_ *f*_*m,y*_ *f*_*m,z*_]^T^ measured at the contact point and the CR’s Kalman-estimated wrist position *p*_*KF*_ = [*p*_*KF,x*_ *p*_*KF,y*_ *p*_*KF,z*_]^T^ were used to define the desired position of the robot end-effector as *P*_*x*_ = *P*_*KF,x*_ + *k*_*f*_ *(f*_*m,x*_ *– f*_*des*_) and *p*_*z*_ = *p*_*KF,z*_ + *k*_*f*_*(f*_*m,z*_ – *f*_*des*_). The desired contact force *f*_*des*_ was set to 0.3 N and the gain *k*_*f*_ was set to 0.00004 m/N, thus regulating the robot motion at the speed of 2.5 mm/s for *f*_*m,i*_ – *f*_*des*_ = 1*N* at the 500 Hz update cycle. For the AP direction, the desired robot position was *P*_*y*_ = *P*_*KF,y*_. Roughly speaking, the presented controller was adding a delta of position *k*_*f*_(*f*_*m*_ – *f*_*des*_) to ML and vertical directions if the measured force was different than *f*_*des*_ = 0.3N. If the measured force was larger than 0.3 N, the delta of position was negative and the robot moves slightly back to reduce the force. If the measured force was smaller than 0.3 N, the delta of the position was positive and the robot pushed slightly against CR’s wrist to remain in contact. In this way, the end-effector kept in contact with the user’s wrist while maintaining low interaction forces. The forces were not different between the two rIPT modes. As expected, the average contact force was close to the prespecified value of 0.3N (average force=0.32N, SD 0.09).

## Results

Table 1 summarizes all statistical comparisons. The MFR amplitude in the horizontal plane was not affected by the IPT condition. All three IPT conditions resulted in comparable amplitudes (hIPT: mean=35.8 cm, SD 1.5; rIPTanticip: mean=35.4 cm, SD 1.4; rIPTfollow: mean=35.1 cm, SD 1.5). Average (Fig. 4a) and peak planar reaching velocity (Fig. 4b) were slower in both rIPT conditions compared to hIPT. The directional angle of reaching in the horizontal plane was more straight ahead in the rIPTfollow condition (AV angle=-0.83 deg, SEM 1.84) and a tendency of less lateral drift in rIPTanticip (AV angle=-1.34 deg, SD 1.92) compared to hIPT (AV angle=-4.55 deg, SD 2.12). Orthogonal deviation from a straight line, in terms of both the average (Fig. 4c) and summed deviation (Fig. 4d) as well as the variability, was lower in hIPT than rIPTanticip. Path length was not altered by the IPT conditions but the normalized path length indicated less curvature in rIPTfollow compared to rIPTanticip (Fig. 4e).

**Table 1.**
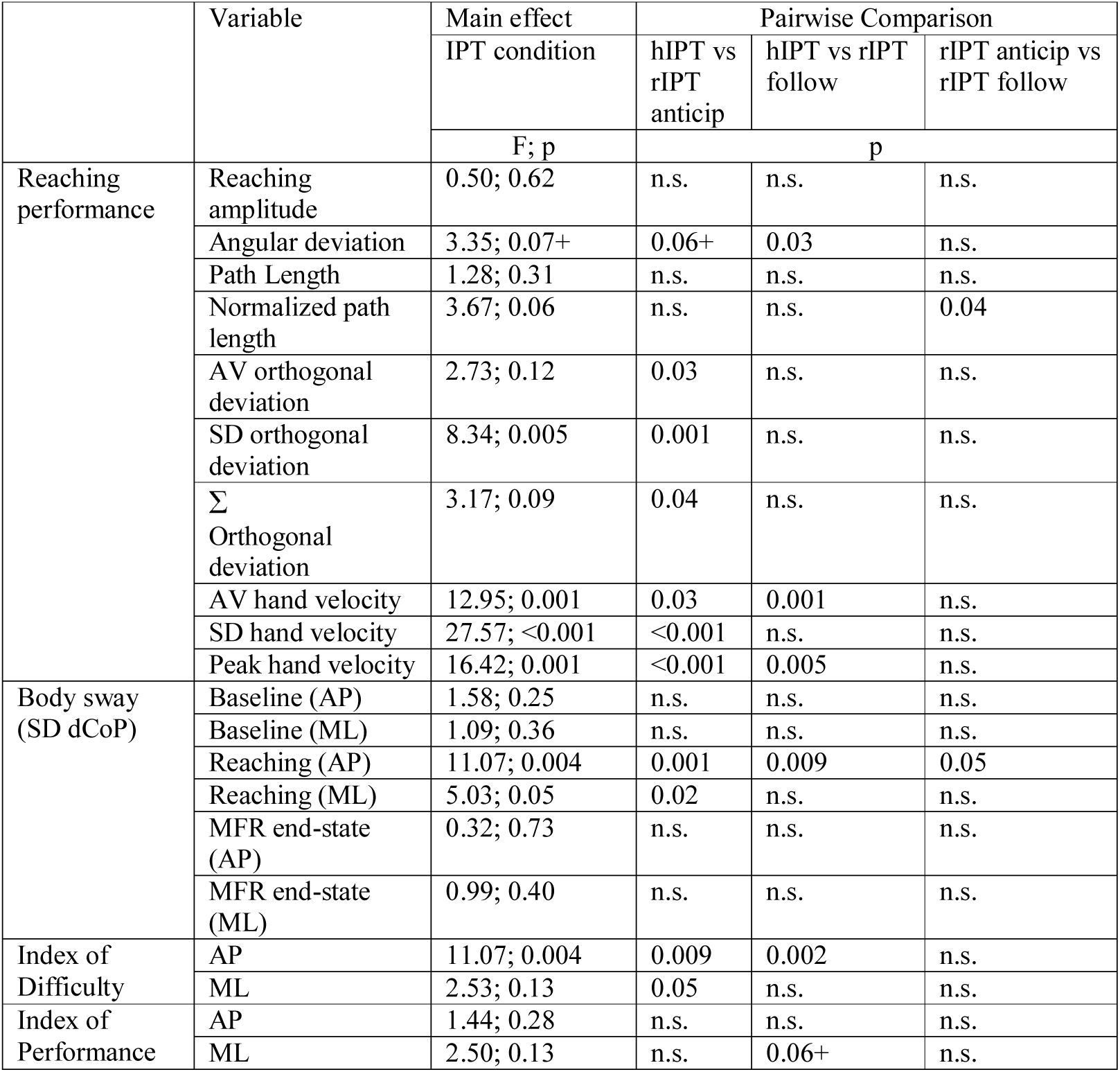
Summary of all statistical comparisons. IPT: interpersonal touch; hIPT: human IPT; rIPT anticip: robotic IPT anticipating; rIPT follow: robotic IPT following; SD dCoP: standard deviation of Centre-of-Pressure velocity; AP: anteroposterior; ML: mediolateral; MFR: maximum forward reach. +: marginally significant; n.s.: not significant.

**Figure 4.**
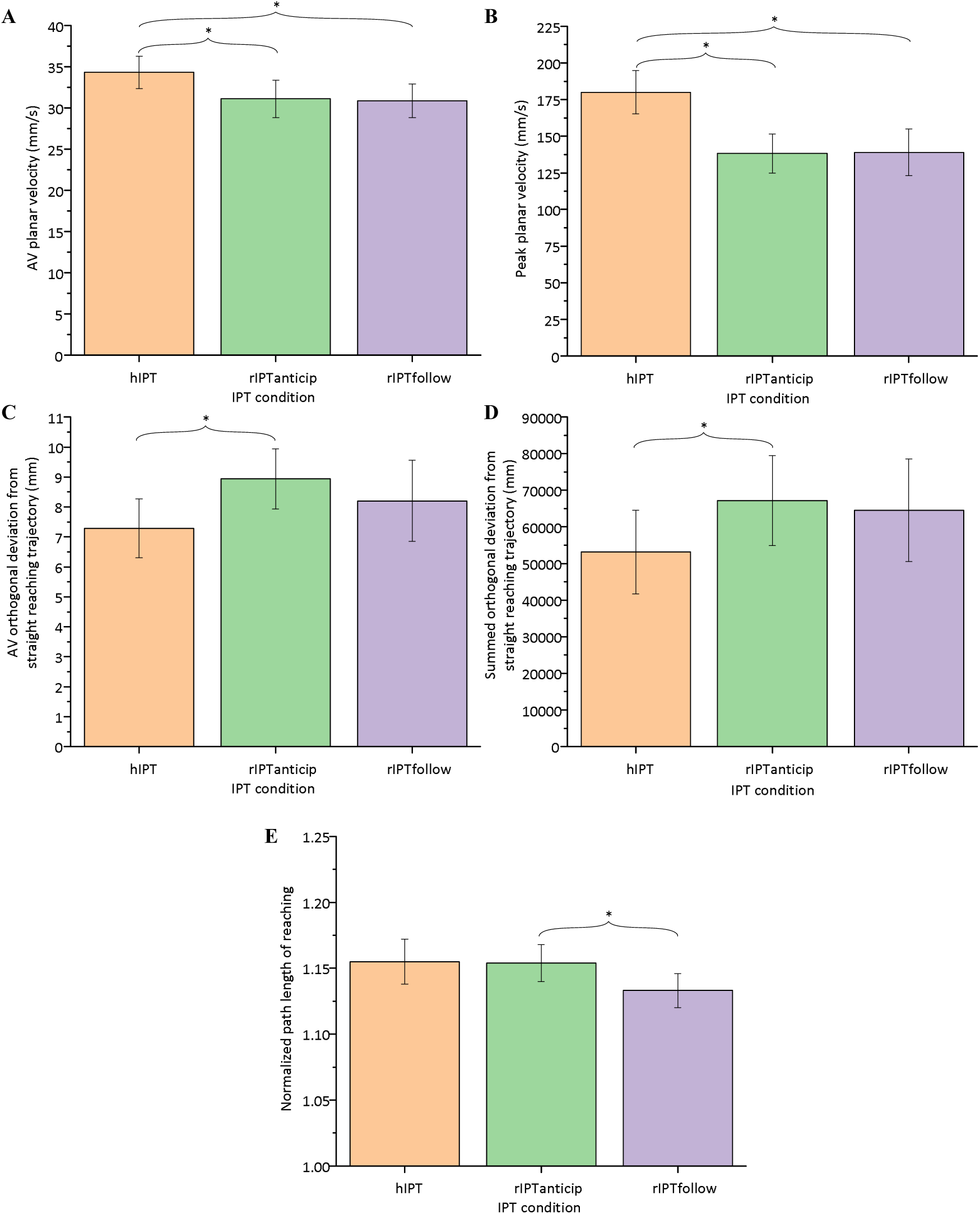
Parameters of reaching performance as a function of the interpersonal touch (IPT) condition: (A) average planar velocity, (B) peak planar velocity, (C) average orthogonal deviation from a straight line linking the start to the end positions, (D) summed orthogonal deviation from a straight line, (E) normalized path length of reaching. Error bars show the standard error of the mean across participants. Horizontal brackets indicate significant within-subject post-hoc single comparisons (p<0.05). hIPT: human IPT; rIPTfollow: robotic IPT in follower mode; rIPTanticip: anticipatory robotic IPT.

Sway variability in either the AP or ML directions was not different between the three IPT conditions in the baseline phase and the MFR end-state. During the reaching phase, however, AP sway variability was reduced in both conditions involving rIPT compared to hIPT (Fig. 5a) and rIPTanticip compared to rIPTfollow. In contrast, only rIPTanticip showed reduced ML sway variability compared to hIPT (Fig. 5b).

**Figure 5.**
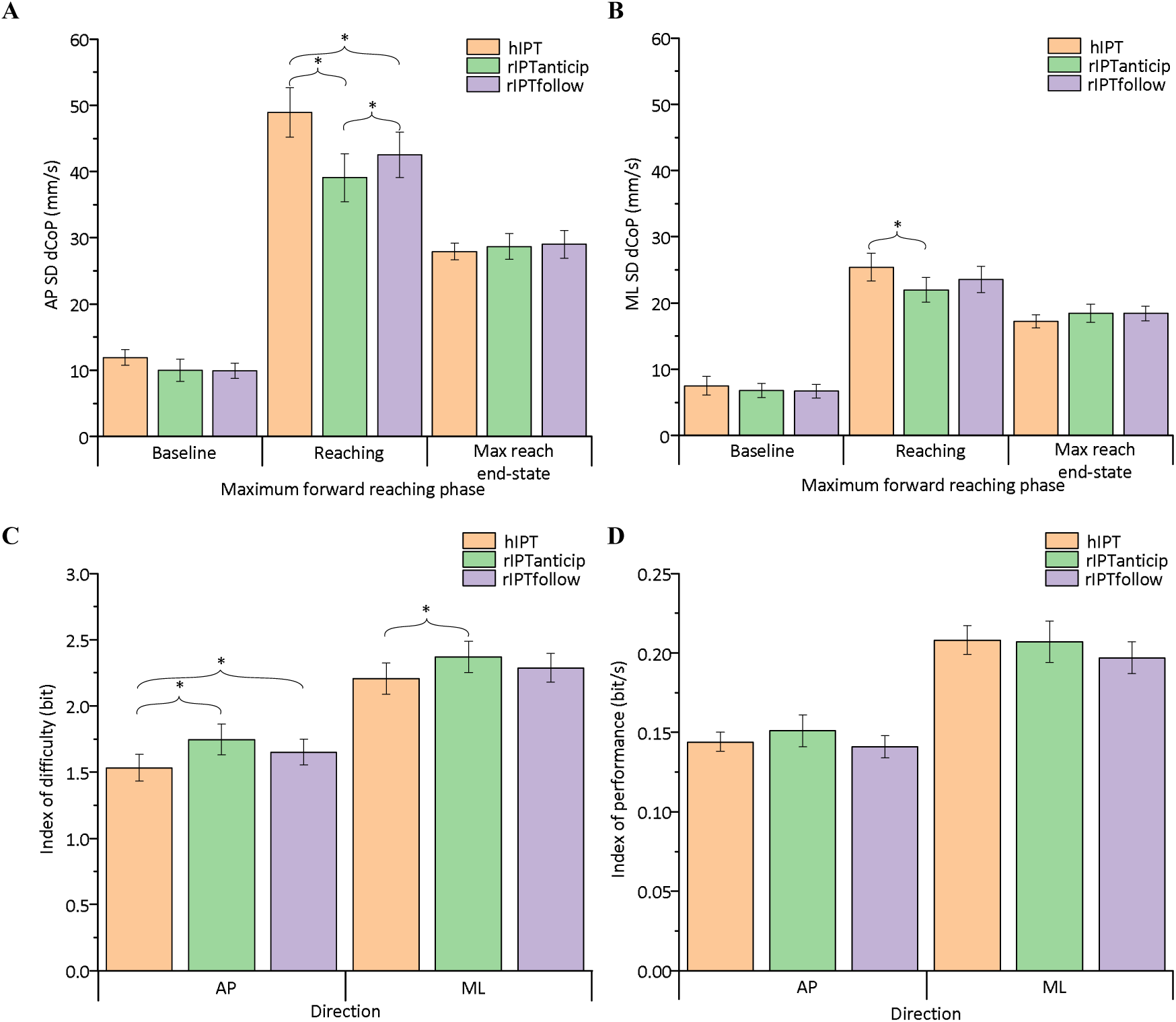
Body sway in terms of the standard deviation of Centre-of-Pressure velocity (SD dCoP) as a function of the interpersonal touch (IPT) condition in the anterior-posterior (A) and mediolateral (B) direction in all three phases of the Maximum Forward Reaching (MFR) task. Index of difficulty (C) and Index of performance (D) in each IPT condition for both directions. Error bars show the standard error of the mean across participants. Full horizontal brackets indicate significant within-subject post-hoc single comparisons (p<0.05). hIPT: human IPT; rIPTfollow: robotic IPT in follower mode; rIPTanticip: anticipatory robotic IPT.

The IoD differed between the three conditions in the AP direction., with the lowest scores in hIPT compared to both rIPT conditions. In the ML direction, hIPT had a lower IoD score compared to rIPTanticip only (Fig. 5c). In contrast, no difference in the informational “throughput” (IoP) was observed between the three conditions (Fig. 5d).

## Discussion

Our study contrasted the effects of deliberately light interpersonal touch received by a robotic system on the control of movements and body balance during maximum forward reaching in healthy young adults. Changes in spontaneous MFR behaviour and body sway were assessed as a function of the robotic system’s mode of control (follower vs anticipation) with respect to CR’s movements. With respect to the body sway in the MFR baseline or end-state as well as the achieved MFR amplitude, rIPT was as efficient as hIPT. The observed changes in reaching behaviour with rIPT coincided with reductions in body sway during the reaching phase in the same condition: rIPTanticip provided the best stabilization of all three IPT conditions. The Index of Difficulty indicated increased behavioural difficulty in the two robotic conditions compared to hIPT, despite the fact that the Index of Performance indicated similar informational throughput between the three conditions. On a qualitative level, however, rIPTfollow resulted in intermediate behavioural alterations, less different to hIPT than rIPTanticip. This observation might imply that in hIPT the human contact provider coordinated the movements in a reactive fashion as well, potentially in follower mode due to visual dominance or as the more optimal strategy due to the inability to stem the computational complexity of predicting CR’s trajectory.

In our current study, the provision of IPT by the CP involved visual feedback of CR and his or her movements.As this would be more similar to the optical tracking of CR’s motion used by the robotic system. In human pairs, the presence of visual feedback with habitual visual dominance is likely to turn the CP into a follower of CR’s movement (Steinl & Johannsen, 2017). Assessing HHI as well as HRI in a single degree of freedom object manipulation task, Groten et al. (Groten, Feth, Goshy, et al., 2009; Groten, Feth, Klatzky, Peer, & Buss, 2009) characterized inter-agent dominance as a function of the interaction force with dominance varying flexibly between both partners in a joint action. Generally speaking, in most physical interactions between two human individuals leader-follower relationships are not necessarily fixed. It seems to be the case, however, that the more adaptive individual, for example the person on whom fewer requirements to fulfill specific movement contraints are imposed, is more likely to take a follower role (Skewes, Skewes, Michael, & Konvalinka, 2015).

Despite impressive advances in the recent decade, current robotics engineering is still distant from developing robotic systems able to assist human individudals socially, especially during postural activities and balance exercises (Sheridan, 2016). In the both rIPT conditions of the current study, the dynamics of the robotic system were not independent but in one way or another a direct consequence of CR’s movements. Despite the lack of any real “social cognitive” capabilities of the robotic system, this fact can nevetheless be interpreted as highly precise responsiveness, which a real human CP could never match. We assume that participants were not able to consciously preceive any difference between the anticipatory and follower rIPT modes, just an absolute timing difference of 20 ms, and therefore would not change their behaviour voluntarily. Possibly due to a shift in participants from less to more reactive, feedback-dependent postural control, CRs reduced their reaching velocity to adjust their movements more precisely to the current position of the robotic end-effector and for the same to stay in contact with their wrist. These concerns could have been even more prominent in the rIPTanticip condition than in rIPTfollow.

### Reaching performance and body sway

An increased MFR amplitude would demonstrate improved confidence in the ability of keeping own body balance stable while approaching one’s forward limits of stability (Duncan, Weiner, Chandler, & Studenski, 1990; Maki & McIlroy, 2006). As we did not observe any differene in reaching amplitude between all three forms of IPT, it means that IPT provided by a robotic system does not disrupt or distract the human CR. During the reaching phase, the facilitation of stabilization of body sway by rIPT tended to surpass the effect of hIPT, especially in a robotic control mode involving anticipation. This shows that rIPT does not destabilize CR’s postural behaviour but can lead to a further reductions in behavioural variability. Nevertheless, human CRs altered their MFR behaviour when IPT was provided not by the human partner but by the robotic system. The most obvious changes were general reductions in the average and peak planar MFR velocity with rIPT. As body sway tended to be reduced in these situations, these adjustments to the robotic CP could reflect a trade-off between speed and accuracy [Fitts, 1954]. According to this interpretation, participants may have effectively controlled sway variability in order to meet any perceived difficulty increase in rIPT resulting from “hardware” constraints imposed by technical limitations of the robotic system and “soft” constraints in terms of fulfilling the task goal of MFR with rIPT support (Bardy, Marin, Stoffregen, & Bootsma, 1999; Scholz & Schoner, 1999). “Asssist-as-needed” (Cai et al., 2006) robotic devices will provide corrective forces only if a participant’s limb movement kinematics hit the walls of a predefined “virtual tunnel” (Duschau-Wicke, von Zitzewitz, Caprez, Luenenburger, & Riener, 2010). These “patient-cooperative” robotic assistive devices may improve the outcome of gait training but also body balance in stroke rehabilitation (Srivastava et al., 2015; Srivastava et al., 2016). Assist-as-needed robotic approaches translate into corrective forces keeping an individual’s body or limbs within an initially defined “normal” range. In contrast to such “positive” force feedback, in which a robotic system aims to guide a participant’s limb along a specific trajectory by applying a corrective force, our deliberately light interpersonal touch paradigm could be described to act with “negative” force feedback. This means that if participants stray from a reaching trajectory, they will perceive a momentary reduction in touch, which might cue them to perform a subtle correction with the intention to minimize contact force variability. The robotic system in our study was controlled according to this principle, and we believe it imitated CR’s behaviour more naturally. At the same time, the reaching trajectory was not prespecified within the robotic system but emerged as a compromise between the CR and the respective CP. In this sense, the CR’s movement range remains completely unconstrained. Any constraints result from the “social” context of the HHI or HRI system.

### Human-robotic movement coordination

Haptic interactions between caregiver and patient play an prominent role in cooperative and collaborative human-human sensorimotor interactions in physical rehabilitation (Sawers & Ting, 2014). More recently, Haarman et al. (Haarman et al., 2017) investigated the balance-assistive forces applied by therapists to the pelvis of patients during gait training. Using force-torques sensors, they quantified the predominant corrective forces applied by the therapists in the mediolateral direction to both sides of the hips at about 9N, amounting to approximately 2% of participants’ body weight. Compared to the forces imposed by the robotic systems in our current study, the forces applied by the therapists are still magnitudes greater.

In a cooperative physical HHI, the relationship between interaction forces and movement kinematics is important for communicating intended movement direction (Mojtahedi, Whitsell, Artemiadis, & Santello, 2017; Sawers et al., 2017; Takagi, Usai, Ganesh, Sanguineti, & Burdet, 2018). Gentry and Murray-Smith (Gentry & Murray-Smith, 2003) described the influence of haptic signals used for coordination and synchronization in human dancing. Hoelldampf et al. (Hoelldampf, Peer, & Buss, 2010) used interaction forces to adjust and optimize the robot’s motion in a system designed for human-robot interactive dancing. Similarly, Chen et al. (Chen et al., 2017; Chen et al., 2015) developed a mobile robotic system responsive to interaction forces to practice dance stepping with a human partner. Response gain and compliance of the robot’s effectors altered human upper body posture and human-robot coordination. Interestingly, the majority of human partners perceived the robots as following their movements (Chen et al., 2015).

In this context it is remarkable that rIPTfollow led to the straightest forward reaching trajectories with least amount of medial drift. This could mean that a robotic system that emphasizes a reactive follower strategy is a better haptic “communicator” in the sense that it made participants to “listen” more closely to the haptic feedback they received. Possibly, participants interpreted rIPT as more reliable as a relative spatial reference and therefore adjusted their reaching movements more in a feedback-driven manner. In contrast, although rIPTanticip also tended towards a more straight ahead reaching movement, the condition showed the greatest and most variable orthogonal deviation from a straight line connecting the start and end point. The robotic system in leader mode could have actually “misguided” participants in the sense, that it tried to anticipate a participant’s next position and so reinforced a participants’ tendency to deviate from their current trajectory. That this interaction did not cause excessive deviations of the reaching trajectory could be a result of the tighter bounds applied to variability of body sway in rIPTanticip.

Mohan et al. (Mohan, Mendonca, & Johnson, 2017) assessed the interactions between a therapist and a stroke patient in the less complex situation of raising and drinking from a cup with the assistance of the therapist. By analyzing the both partners’ movement kinematics, they concluded that the strength of the interpersonal coupling varied as a function of the task’s phase with stronger interaction at the beginning and the end of the action (Mohan et al., 2017). In our current study, the robotic system operated in a single control mode throughout an entire trial. In terms of shaping the participants’ MFR behaviour it might be even more optimal, if the robotic system had switched from a leader mode in the baseline phase and the end-state to a follower mode during the reaching phase.

## Conclusions

Beneficial deliberately light interpersonal touch for balance support during maximum forward reaching is easily provided by a robotic system even when it is mechanically uncoupled to the human contact reveicer. This effect does not rely on the system’s capability to predict the future position of the contact receiver’s wrist. The effects the uncoupled robotic IPT in reactive following mode were comparable to human IPT on most parameters. As the robotic system itself was not designed for any form of “social” cognition or explicit haptic communication, our study nevertheless demonstrates that robotic IPT can be used to implicitly “nudge” human contact receivers to alter their postural strategy for adapting to the robotic system without any decrements in their postural performance during maximum forward reaching.

## Acknowledgements

We thank Prof. G. Cheng, Prof. M. Buss, Prof. S. Hirche, Dr. K. Ramirez–Amaro, and J. R. Guadarrama Olvera for providing the experimental infrastructure and S. M. Steinl and M. Langer for their involvement in data acquisition. Funding: we acknowledge the financial support by the federal Ministry of Education and Research of Germany (BMBF; 01EO1401), the Deutsche Forschungsgemeinschaft (DFG) through the TUM International Graduate School of Science & Engineering (IGSSE), German Academic Exchange Service (DAAD), and Helmholtz Associate.

## Author contributions

K.P. analyzed datasets for the experiment, interpreted results, and wrote the paper. S.E., and H.S. designed and performed the experiment, interpreted results, and wrote the paper. M.S. designed and performed the experiment, interpreted results, and wrote the paper. L.J. and D.L. designed the experiment, interpreted results, and wrote the paper.

## Key points

1. Robotic light touch supports human balancing performance
2. Human participants adapt to the specific affordances of robotic light touch support
3. Subtle differences in the robotic modes of interaction have behavioural effects on the human performer

